# Selective medial septum lesions in healthy rats induce longitudinal changes in microstructure of limbic regions, behavioral alterations, and increased susceptibility to status epilepticus

**DOI:** 10.1101/2023.08.11.553036

**Authors:** Hiram Luna-Munguia, Deisy Gasca-Martinez, Alejandra Garay-Cortes, Daniela Coutiño, Mirelta Regalado, Ericka de los Rios, Paulina Villaseñor, Fernando Hidalgo-Flores, Karen Flores-Guapo, Brandon Yair Benito, Luis Concha

## Abstract

Septo-hippocampal pathway is crucial for physiological functions and is involved in epilepsy. Its clinical monitoring during epileptogenesis is complicated. We aim to evaluate tissue changes after lesioning the medial septum of normal rats and assess how the depletion of specific neuronal populations alters the animals’ behavior and susceptibility to establishing a pilocarpine-induced *status epilepticus*. A total of 64 young-adult male Sprague-Dawley rats were injected into the medial septum with vehicle or saporins (GAT1 or 192-IgG for GABAergic or cholinergic depletion, respectively; n=16 per group). Thirty-two animals were used for diffusion tensor imaging (DTI); they were scanned before surgery and 14 and 49 days post-injection. Fractional anisotropy and apparent diffusion coefficient were evaluated in the fimbria, dorsal hippocampus, ventral hippocampus, dorso-medial thalamus and amygdala. Between scans 2 and 3, animals were submitted to the elevated plus-maze, open-field test, rotarod test, Y-maze and water-maze. Timm, toluidine and Nissl staining were used to analyze tissue alterations. Twenty-four different animals received pilocarpine to evaluate the latency and severity of the *status epilepticus* two weeks after surgery. Eight animals were only used to evaluate the extent of neuronal damage inflicted on the medial septum one week after the molecular surgery. Progressive changes in DTI parameters in both the white and gray matter structures of the four evaluated groups were observed. Behaviorally, the GAT1-saporin injection impacted spatial memory formation, while 192-IgG-saporin triggered anxiety-like behaviors. Histologically, the GABAergic toxin also induced aberrant mossy fiber sprouting, tissue damage and neuronal death. Regarding the pilocarpine-induced *status epilepticus*, this agent provoked an increased mortality rate. Selective septo-hippocampal modulation impacts the integrity of limbic regions crucial for certain behavioral skills and could represent a precursor for epilepsy development.

**Significance statement:** The medial septum is believed to be involved in epilepsy. However, whether and how defects in the integrity of each neuronal subpopulation conforming this structure affect gray and white matter structures remains unclear. Here we examine whether the injection of vehicle or partially-selective saporins into medial septum of normal rats play a role in the integrity of specific brain regions relevant to memory formation, anxiety-like behaviors, and susceptibility to *status epilepticus* induction and survival. We find that lesioning the medial septum GABAergic or cholinergic neurons can represent a precursor for behavioral deficits or epilepsy development. Therefore, these results strongly support the idea that modulation of medial septum can be a potential target to improve cognition or reduce seizure frequency.

## 1. Introduction

The medial septum is a small gray matter nucleus located in the middle of the basal forebrain. It contains interconnected cholinergic, GABAergic and glutamatergic neurons that mainly project through the fimbria/fornix to both hippocampal formations. The circuit is completed after some hippocampal GABAergic axons arise from non-principal neurons and extend toward the medial septum (Kiss et al., 1990; Toth et al., 1993; Jinno and Kosaka, 2002; Colom et al., 2005). Several rodent studies applying specific approaches have established that the integrity of each neuronal subpopulation forming this septo-hippocampal network is crucial for certain learning and memory tasks (Walsh et al., 1996; Janis et al., 1998; Johnson et al., 2002; Pang et al., 2011; Roland et al., 2014a, 2014b; Khakpai et al., 2016). Other studies have focused on the role of medial septum cholinergic efferents in anxiety-like behaviors (Lamprea et al., 2000; Pizzo et al., 2002; Zarrindast et al., 2013; Zhang et al., 2017).

Rodent electrophysiological studies have also described the relevance of the septo-hippocampal pathway in generating and modulating the hippocampal theta rhythm (4-12 Hz) (Buzsáki, 2002; Hangya et al., 2009; Vandecasteele et al., 2014; Colgin, 2016; Robinson et al., 2016), a distinctive oscillatory activity that occurs during rapid eye movement sleep, active spatial navigation, and memory processing (Buzsáki, 2002, 2005; Buzsáki and Moser, 2013). However, a reduction of hippocampal theta power not only induces cognitive deficits, it has also been related to epileptic activity in experimental temporal lobe epilepsy (Colom et al., 2006; Chauviere et al., 2009; Lee et al., 2017). Some studies have associated this alteration with a direct GABAergic medial septum lesion (Butuzova and Kitchigina, 2008). Others argue that removing the medial septum cholinergic projections facilitates the initial stages of hippocampal kindling (Ferencz et al., 2001).

Temporal lobe epilepsy (TLE) is the most common type of focal epilepsy in humans (Engel Jr, 1996). Clinically, magnetic resonance diffusion tensor imaging (DTI) has been extensively used as a noninvasive technique to identify subtly disturbed white matter microstructure, detecting abnormalities in several white matter structures (Arfanakis et al., 2002; Concha et al., 2005a, 2010; Gross et al., 2006; Schoene-Bake et al., 2009; Hatton et al., 2020). The fornix, in particular, has shown bilaterally symmetric diffusion abnormalities that correspond to reduced axonal density and myelin alterations (Concha et al., 2010). Notwithstanding the degeneration of hippocampal efferents that directly affect fornix diffusion metrics, the bilaterality of abnormalities of the fornix (in spite of clearly lateralized hippocampal damage) is suggestive of additional anomalies of septo-hippocampal fibers. Brain injury, tumors, and structural malformations are common etiologies of TLE. However, some patients lack any evidence of an initial precipitating injury (Harvey et al., 1997; Sztriha et al., 2002). In such cases, the mechanisms underlying the source of the neurological disorder and the progressive cerebral microstructural changes before the first seizure arises are not yet fully elucidated. Clinically, this analysis is complicated, hence the relevance of animal models. Previous longitudinal studies have described the structural changes in gray and white matter microstructure after experimental *status epilepticus* induction in diverse animal models of TLE (Laitinen et al., 2010; Parekh et al., 2010; van Eijsden et al., 2011; Janz et al., 2017; Salo et al., 2017; Wang et al., 2017; Luna-Munguia et al., 2021). Here, *status epilepticus* is the severe triggering factor that allows monitoring distinct behavioral and neurophysiological features of the TLE pathogenesis. However, there are pending studies that track microstructural and histological changes after a subtle limbic-induced defect that ultimately represent a predisposing factor able to increase seizure susceptibility.

The aim of this study is to investigate the influence of GABAergic or cholinergic medial septum lesions on the temporal changes in gray and white matter microstructure through DTI. We used the immunotoxin GAT1-saporin and the neurotoxin 192-IgG-saporin to selectively target GABA or choline neuron populations of the medial septum, respectively. Then, we evaluated the relevance of each neuronal subpopulation by directly comparing the effects of the lesions on diverse behavioral performances and on seizure development in the pilocarpine-induced *status epilepticus* model. Imaging findings were compared to histological assessments of neurodegeneration and mossy fiber sprouting in specific staining preparations from the same animals.

## 2. Methods

### 2.1 Animals

A total of 64 male Sprague-Dawley rats were provided by our animal facility and used for different experimental purposes (Figure 1). All were individually housed and taken to the assigned room one week prior to the initiation of any experiment. The room had constant controlled conditions (12-h light/dark cycle, 20-22°C, 50-60% relative humidity), and animals had *ad libitum* access to food and water. Every procedure adhered to the guidelines established by our Institutional Ethics Committee for Animal Use and Care (Protocol #105A) following the Official Mexican Standard NOM-062-ZOO-1999/SAGARPA.

**Figure 1.**
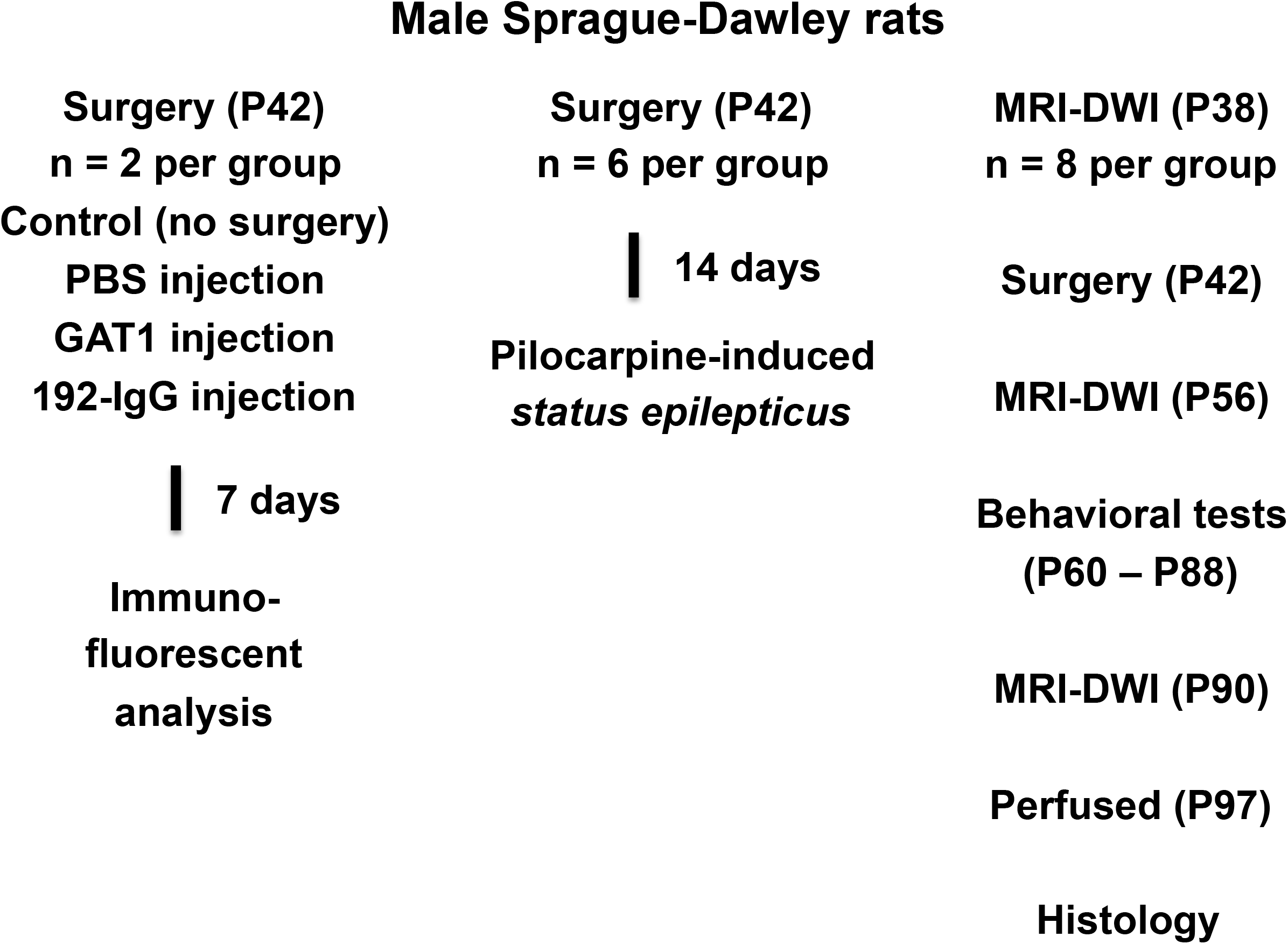
Diagram of the experimental protocol. A group of rats was submitted to the immunofluorescent analysis to evaluate the extent of neuronal damage inflicted on the medial septum after seven days of the injection. The second group of animals was only used to evaluate the latency to *status epilepticus* and the mortality rate. The third group was subjected to MR-DWI scans, behavioral tests, and histological analysis. P: postnatal day.

### 2.2 *In vivo* Magnetic Resonance Imaging (MRI)

Acquisition protocols were carried out at the National Laboratory for MRI using a 7T scanner (Bruker Pharmascan 70/16US). *In vivo* DTI and T2 images were acquired using a 72 mm inner-diameter volume coil for transmission and a 2×2 rat head array coil for reception (Bruker, Ettlingen, Germany).

Thirty-two rats were scanned at three time points: 1) before surgery (postnatal day 38 (P38); see below), 2) two weeks after surgery (P56), and 3) once the animals had finished the five behavioral evaluations (P90; see below). Animals were anesthetized with a 4% isoflurane/O _2_ mixture and a 2% mix to maintain anesthesia during image acquisition. Body temperature was maintained by recirculating warm water underneath the imaging bed, and vital signs were continuously monitored. Anesthesia was discontinued upon completion of the imaging session. Once the animals were fully recovered, they were transferred back to the housing room. Data sets for DTI were acquired using an echo-planar imaging sequence with the following parameters: TR = 2250 ms, TE = 31.32 ms, number of averages = 4, slice thickness = 750 μm, FOV = 20 × 14 mm^2^, matrix = 150 × 104, yielding in-plane resolution = 133 × 135 μm ^2^, scan time = 27 min. Diffusion-weighted images were acquired in forty unique directions (δ/Δ = 2.9/8.7 ms), each with two different b values (650 and 2000s/mm ^2^) along with six images without diffusion weighting. Immediately afterwards, a 5-min fast low-angle shot (FLASH) scan was acquired with the following parameters: TR = 350 ms, TE = 2.4 ms, NA = 3, flip angle = 30°, FOV = 25 × 18 mm^2^, matrix = 250 × 180 (100 μm in-plane resolution), slice thickness = 800 μm, number of slices = 20. The total scan time was 32 minutes.

The rest of the total number of requested animals was only submitted to the 5-min anatomical scan at P38. The aim of this imaging sequence was to detect unexpected brain injury or ventriculomegaly (Luna-Munguia et al., 2020) in the animals assigned to the immunofluorescent or pilocarpine protocols (see below). Animals displaying such abnormalities were discarded from the study.

### 2.3 Image data processing

Diffusion-weighted imaging (DWI) data sets were denoised (Veraart et al., 2016), and linear transformations (12 dof) were used to correct for motion and eddy current-induced distortions. The diffusion tensor from which we obtained the corresponding eigenvalues (λ_1_, λ_2_, and λ_3_) was estimated using the MRtrix 3.0 software package (Tournier et al., 2019). From these, we created quantitative maps of fractional anisotropy (FA) and apparent diffusion coefficient (ADC) (Basser and Pierpaoli, 1996). Regions of interest (ROIs) were manually drawn by three independent reviewers, outlining the dorsal hippocampus, ventral hippocampus, fimbria, dorso-medial thalamus, and amygdala for the three time points. The ROIs were outlined on the color-coded maps of the principal diffusivity (aided by the non-diffusion weighted images). For each structure and animal, diffusion metrics were averaged for all voxels included in the corresponding ROI.

### 2.4 Molecular surgery

Neurons in the medial septum were selectively injured. Rats were randomly divided into four groups according to the medial septum injected agent: GAT1-saporin (325 ng/ µl dissolved in sterile 0.1X phosphate buffered saline (PBS; Sigma-Aldrich); n=16; Advanced Targeting Systems), 192-IgG-saporin (375 ng/µl dissolved in sterile 0.1X PBS; n=16; Advanced Targeting Systems), PBS (0.1X dissolved in sterile saline solution; n=16), and control (no surgery; n=16). After ketamine/xylazine mixture anesthesia (70 mg/kg and 10 mg/kg, respectively; administered intraperitoneally), rats were placed in a Stoelting stereotactic frame. An incision was made midline on the scalp, and the skull was exposed using sterile surgical techniques. Based on preliminary studies designed to determine the medial septum coordinates in P42 animals, we drilled a small hole into the skull at the following coordinates: anteroposterior +0.1 mm, lateral -2 mm, ventral from skull surface 6.8 mm. Then, a 5 µl Hamilton syringe was adapted to the stereotactic frame arm and it was angled up to 20°. The syringe was gradually lowered and infused 1.2 µl of the corresponding agent at a rate of 0.1 µl/min using a WPI Microsyringe Pump Controller. The syringe was left in place for 5 min after injection to ensure agent diffusion. Then, the syringe was gradually removed to avoid liquid suction. The wound was sutured and animals received ketorolac (30 mg/ml im; PiSA) every 24 h for two days. Thirty-two animals were randomly transferred back to the housing room located in the Institute’s animal facility building for posterior pilocarpine experiments (n=24; see below) or immunofluorescent analysis (n=8; see below). The other 32 rats were taken to a new experimentation room (same conditions except for the inverted light/dark cycle) located in the Institute’s Behavioral Analysis Unit building.

### 2.5 Immunofluorescent analysis

The extent of neuronal damage inflicted on the medial septum was evaluated. Seven days after surgery, two animals per group were intracardially perfused with 0.9% NaCl solution followed by 4% paraformaldehyde (PFA; Sigma-Aldrich) solution. Brains were removed and post-fixed in fresh 4% PFA solution for 24 h. Then, they were immersed until sinking in 20% and 30% sucrose solution. Specimens were carefully frozen using dry ice and stored at - 80°C. Twenty micron-thick coronal slices encompassing the medial septum were cut in a cryostat (Leica Biosystems 3050S) and stored in cold 1X PBS until the immunofluorescence staining. Brain sections were first incubated in blocking solution (2% bovine serum albumin [Sigma-Aldrich] and 0.3% triton X-100 [ThermoFisher] in 1X PBS) for 45 min at constant agitation and 4°C. Then, for double immunofluorescence stainings, four slices per brain were used to determine the loss of GABAergic neurons or cholinergic neurons. For the former, sections were incubated with the primary antibodies anti-parvalbumin mouse monoclonal (1:300; Sigma-Aldrich) and anti-NeuN rabbit polyclonal (1:500; Abcam) for 24 h at 4°C, then rinsed three times for 10 min in 1X PBS and incubated for 4 h at 4°C with the fluorescently tagged secondary antibodies (Invitrogen; AlexaFluor 647 goat anti-mouse and AlexaFluor 488 donkey anti-rabbit; diluted at 1:400 and 1:500, respectively, in blocking solution). Finally, sections were rinsed three times for 10 min in 1X PBS and cover-slipped with microscope cover glass. Images were collected using a laser confocal microscope (Zeiss LSM 780 DUO). To determine the loss of cholinergic neurons, slices were incubated with the primary antibodies anti-choline acetyltransferase (ChAT) rabbit polyclonal (1:500; Sigma-Aldrich) and anti-NeuN mouse monoclonal (1:500; Abcam) for 24 h at 4°C, then washed and incubated as previously described with the secondary antibodies (Invitrogen; AlexaFluor 555 goat anti-rabbit and AlexaFluor 488 goat anti-mouse; both diluted at 1:500 in blocking solution). Slices were rinsed and cover-slipped as mentioned above. Images were captured using an Apotome-Zeiss fluorescence microscope connected to a computer with AxioVision software. The number of medial septum parvalbumin- or ChAT-positive cells was counted offline using Image J software.

### 2.6 Pilocarpine-induced *status epilepticus*

Two weeks after surgery, six animals per group were submitted to a method based on Luna-Munguia et al. (2017). Here, 56-day-old rats received an injection of pilocarpine hydrochloride (340 mg/kg ip; Sigma-Aldrich) 20 min after being injected with atropine sulfate (5 mg/kg ip; Sigma-Aldrich). Experienced researchers evaluated the animals’ behavioral changes until *status epilepticus* was reached. Those animals that did not develop it within 40 min received an additional intraperitoneal dose of pilocarpine (170 mg/kg). Diazepam (10 mg/kg ip; PiSA) was injected after 90 min of *status epilepticus*. For this study, we only evaluated the latency to *status epilepticus* and the mortality rate. Rats were orally supplemented with Ensure for three days following the induction and used for another research protocol.

### 2.7 Behavioral tests

All behavioral tests were carried out under controlled environmental conditions (temperature [20-22°C], relative humidity [50-60%], and light intensity [dim illumination]). To avoid circadian alterations, all experiments were conducted during the dark/active cycle of the rats (specifically, between 09:00 and 13:00 h) at the Institute’s Behavioral Analysis Unit building. Eighteen days after surgery (or four days post-second MRI scanning), eight animals per group were submitted to a battery of tests as follows:

#### 2.7.1 Elevated plus-maze test

The elevated plus-maze has been described as a simple method to assess anxiety-related behaviors in rodents (Walf and Frye, 2007). It comprises an opaque Plexiglas apparatus of four crossed arms (50 cm long × 10 cm wide) and a central region (10 cm × 10 cm) elevated 50 cm above the ground floor. Two arms were enclosed by 40-cm-high walls, and two arms were open. The elevated plus-maze was placed close to the center of the room, and the level of illumination was approximately 100 lx in the open arms and 35 lx in the closed arms. The animals were habituated to the testing room for at least one hour prior to the evaluation. At the beginning of the experiment, each rat (P60) was individually placed in the maze center, facing an open arm. The animals’ exploratory activity was video recorded for 5 min. The apparatus was thoroughly cleaned after the removal of each animal. By using the SMART 3.0 video tracking software (Panlab Harvard Apparatus, Spain), two parameters were evaluated: 1) the total number of entries into the open arms and 2) the time spent in the open arms. All four paws had to cross the entrance to the open or closed arm to be considered an entry into that arm (Weninger et al., 1999). Twenty-four hours later, animals were submitted to the next test.

#### 2.7.2 Open-field test

Open-field activity measures naturally occurring behaviors exhibited when the animal explores and interacts with its surroundings. This test provides reliable data regarding gross motor function and specific activity related to psychological conditions such as anxiety-like behaviors (Prut and Belzung, 2003). The evaluation was conducted in a room isolated from acoustic interruptions. For assessment of locomotor behavior, we used the SuperFlex/Fusion system in the Digiscan Animal Activity Monitors (Omnitech Electronics, USA). Briefly, each rat was individually placed in a well-lit testing chamber constructed of Plexiglas (40 cm long × 40 cm wide × 30 cm height) for a 24-h observation period. Before recording started, some food pellets were dropped on the floor, and a plastic bottle filled with water was secured in one corner of the chamber. The animals were allowed to freely explore the box, and locomotion activity was captured by three-paired 16-photocell Superflex Sensors, which transmitted the location data to the Superflex Node. The apparatus was thoroughly cleaned after the removal of each animal. Data were processed using Fusion Software (version v5.3), and the following parameters were evaluated: 1) total distance traveled and 2) time spent in the center of the chamber (Miyakawa et al., 2001). The animals were taken to the third test the next day.

#### 2.7.3 Rotarod test

Motor coordination and balance were evaluated using a rotarod apparatus (IITC Inc. Life Science, USA). Testing consisted of four trials each day for three consecutive days (P63 - P65). The rod accelerated constantly and uniformly from 5 to 40 rpm in 60 s. The apparatus was thoroughly cleaned after the removal of each animal. Data analysis was based on the maximal time that the rat was able to stay on the rotating rod (Misumi et al., 2016). Animals were not submitted to any task for the next four days.

#### 2.7.4 Y-maze alternation test

On P70, this alternation task was used to measure spatial working memory during the exploratory activity of the animals (Hughes, 2004). Briefly, this apparatus has three identical arms (arm width, 10 cm; arm length, 94 cm; wall height, 25 cm) oriented 120° from each other. Each rat was placed in the center of the device and allowed to freely explore for 10 min. The sequence and number of arm entries were tracked for each animal. After each trial, the apparatus was cleaned with 70% ethanol to minimize olfactory cues. Arm entry was counted when all four paws were placed inside the arm. To calculate the percent of spontaneous alternation: [(number of alternations)/(total number of arm entries – 2)] × 100. Five days later, animals were submitted to the next test.

#### 2.7.5 Morris water maze test

This test evaluated spatial learning and memory (Morris, 1984) in our P75 animals by using a circular tank (172 cm diameter, 78 cm deep; San Diego Instruments, USA) located in a room isolated from noise where visual extra-maze cues were attached to the walls. The pool was divided into four equal quadrants and filled to 75% of its capacity with water at 25 ± 1°C. Then, inside the pool, a 15-cm-diameter escape platform was placed in the middle of the north-east zone and submerged 2 cm below the water surface (so animals cannot see it). A video camera mounted above the pool was used to record all the swimming trials. Videos were analyzed offline using SMART 3.0 software.

In brief, the protocol was used as follows: rats were trained for three consecutive days (four trials per day and 5 min between trials). On each trial, the animal was placed in the tank (facing the wall) in one of the four quadrants and allowed to escape and seek the hidden platform (it is important to highlight that a different starting point was used in each trial). If an animal did not escape within 60 s, it was manually guided to the escape platform. After climbing the platform, animals were allowed to stay there for 20 s. The latency to find and climb the hidden device was recorded and used as a task acquisition measurement. Retention was tested on the fifth day, removing the escape platform and submitting each animal to a single 1-min trial (named transfer test).

### 2.8 Histological analyses

One week after the last MRI scan was done, animals were overdosed using an intraperitoneal injection of sodium pentobarbital (Aranda). Four animals per group were randomly chosen for toluidine blue staining, two for Timm staining, and two for Nissl staining.

#### 2.8.1 Timm staining

Mossy fiber sprouting in the molecular layer of the dentate gyrus, elucidated by Timm staining, was evaluated in two animals per group. The protocol was partially based on Cintra et al. (1997). Briefly, animals were transcardially perfused with a 0.9% NaCl solution followed by a 0.1 M sodium sulfide nonahydrate (Meyer) solution and a Karnovsky solution (1% PFA, 1.25% glutaraldehyde in 0.1X PBS; pH 7.4). Then, the brains were removed and post-fixed in fresh Karnovsky solution mixed with 30% sucrose solution to cryoprotect the tissue. Once precipitated, each brain was sliced using a cryostat (Leica Biosystems 3050S). Forty-micron-thick coronal sections were obtained, mounted on electro-charged slides, and submerged in Timm staining solution (30% arabic gum, 0.13 M citric acid [Golden Bell], 0.08 M sodium citrate [Golden Bell] buffer [pH 4], 0.15 M hydroquinone [Sigma-Aldrich], 0.005 M silver nitrate [JT Baker]) for 90 min in the dark at room temperature. Finally, staining sections were counterstained by using a 0.1% cresyl violet solution. Images were taken using an AmScope trinocular microscope fitted with a digital camera.

#### 2.8.2 Toluidine blue staining

Rats were intracardially perfused with a 0.9% NaCl solution, followed by a 4% PFA and 50% glutaraldehyde (Electron Microscopic Sciences) solution (1:25; pH 7.4). Once fixed, the entire brain was removed and transferred to a fresh PFA-glutaraldehyde solution and stored for 24 h at 4°C. The next day, each brain was placed in a new container with fresh 4% PFA solution and stored at 4°C until the dissection day, when two tiny tissue blocks spanning the left and right dorsal hippocampus coupled to their respective fimbria were dissected from each brain. Specimens were washed with 0.1 M sodium cacodylate buffer (Electron Microscopic Sciences) and 3% glutaraldehyde solution for 60 min. Then, they were treated with 0.1% osmium tetroxide (Electron Microscopic Sciences) and 0.1 M sodium cacodylate buffer solution for 6 h. Next, each sample was washed twice in fresh 0.1 M sodium cacodylate buffer (10 min each), dehydrated twice with ethyl alcohol (starting at 10%, 20%, 30% until reaching absolute concentration; 10 min each), and treated with oxide propylene (Electron Microscopic Sciences) for 30 min. Tissue was submerged in a mixture of oxide propylene and epoxy resin (Electron Microscopic Sciences, EPON Kit Embed-812) for 48 h. Samples were removed and placed in fresh epoxy resin for 5 h, then taken out and placed in BEEM capsules containing fresh epoxy resin, moved to an oven and left there at 60°C for 36 h. After polimerization, each block was sliced using an ultramicrotome (RMC Power Tone XL). Sections (600 nm thick) were stained with toluidine blue (HYCEL) and 5% sodium tetraborate (CTR Scientific) solution. Images were taken using an AmScope trinocular microscope fitted with a digital camera.

#### 2.8.3 Nissl staining

Animals were intracardially perfused with 0.9% NaCl solution followed by a 4% PFA solution (pH 7.4). Brains were removed and post-fixed in fresh 4% PFA solution for 24 h at 4°C. Then, they were immersed until sinking in 20% and 30% sucrose solution at 4°C. Specimens were carefully frozen using dry ice and stored at -80°C until slicing. Brains were sectioned in the coronal plane (90 μm) using a Leica Biosystems 3050S cryostat. The series of sections were stained with cresyl violet and used to identify the cytoarchitectonic boundaries of the dorsal hippocampus, dorsomedial thalamus and amygdala. Images were taken using an AmScope trinocular microscope fitted with a digital camera. The severity of tissue damage was evaluated.

### 2.9 Statistical analysis

Statistical analyses were conducted using GraphPad Prism8. All data were compiled as mean ± SEM. For imaging data, we averaged the left and right ROI values for each structure since no statistical asymmetries were observed in the metrics after paired *t*-test analysis. Results presented as bar graphs or temporal courses were analyzed by a one-way ANOVA or a multiple comparisons two-way ANOVA, respectively; in both cases, a *post hoc* Tukey test was conducted. In all statistical comparisons, significance was assumed at the level of *p* < 0.05.

## 3. Results

### 3.1 Immunofluorescence

One week after injecting the GAT1-saporin into the medial septum, it preferentially reduced the GABAergic neurons (↓73% and ↓80%) (Figure 2C) rather than the cholinergic ones (↓40% and ↓35%) (Figure 2G). Animals injected with 192-IgG-saporin into the medial septum showed an evident reduction of cholinergic (↓89% and ↓90%) (Figure 2H) rather than GABAergic neurons (↓30% and ↓35%) (Figure 2D).

**Figure 2.**
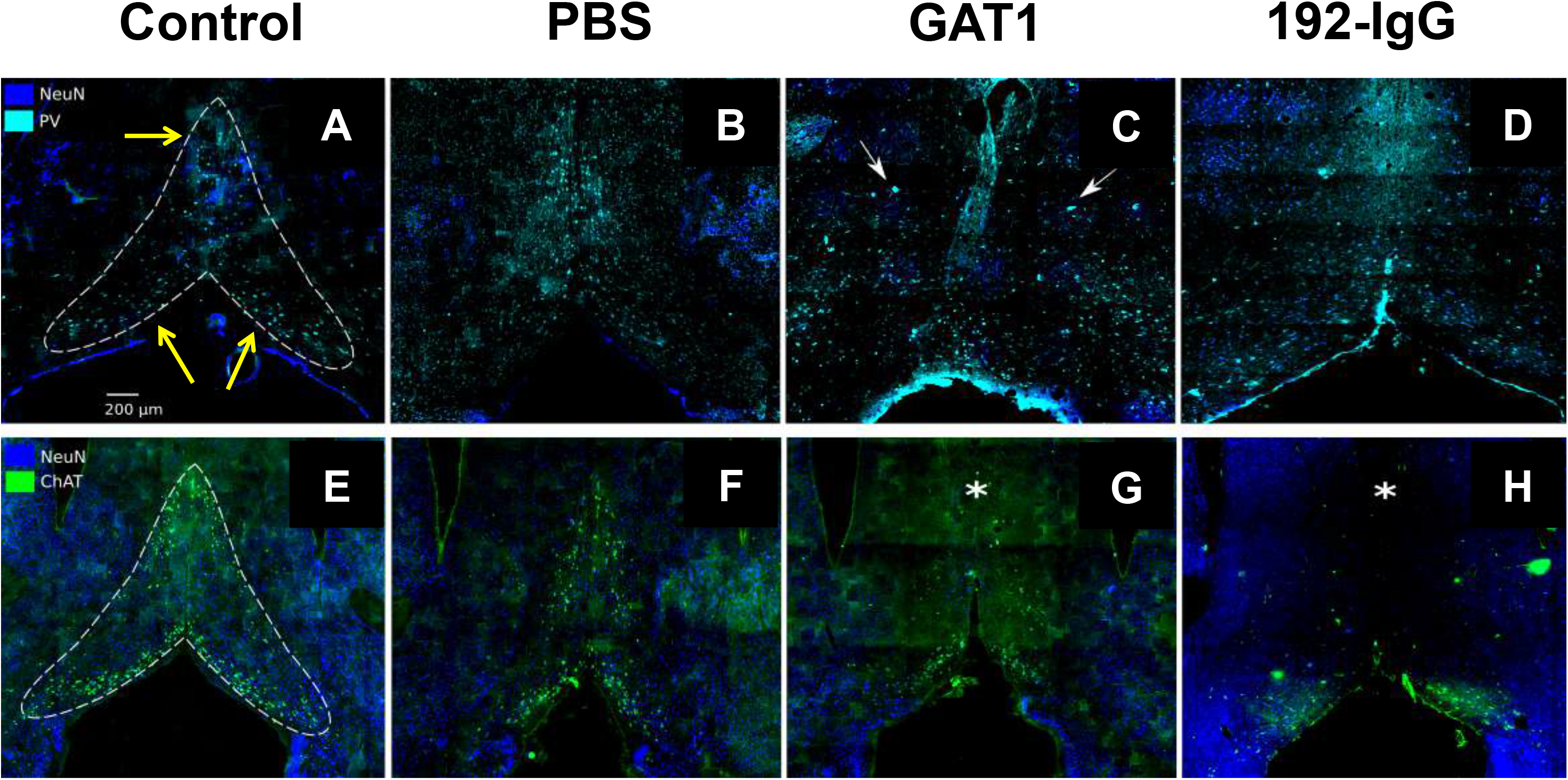
Parvalbumin (PV) and choline acetyltransferase (ChAT) immunopositive neurons, as well as pan-neuronal label NeuN. The dotted lines in **A** and **E** limit the perimeter of an intact medial septum in control rats; yellow arrows indicate the areas where the neuronal quantification was performed. **B** and **F** show a slight reduction of GABAergic or cholinergic neurons, respectively, after medial septum injection of phosphate buffered saline (PBS); **C**: GAT1-saporin injection into the medial septum clearly deformed its structure and mainly reduced the presence of PV-immunopositive neurons (arrows at **C**); GAT1-saporin is partially selective since ChAT-immunoreactive cells are also affected, mainly in the upper part of the medial septum (asterisk) (**G**). A reversed effect occurred with the 192-IgG-saporin injection into the medial septum: ChAT-immunopositive cells were mainly affected (asterisk) (**H**) but PV-immunoreactive cells were also decreased (**D**).

In the case of the PBS-injected rats, a slight decrease of parvalbumin-(↓18% and ↓19%; Figure 2B) and ChAT-immunoreactive cells (↓20% and ↓21%; Figure 2F) was observed when compared to the control subjects (Figure 2A, 2E). This indicates that the mechanical lesion secondary to the injection can induce certain destruction of the GABAergic or cholinergic neuronal populations, albeit not to the extent of the cellular depletion produced by the saporins. After establishing the effectiveness of both saporins, we proceeded to the following experiments.

### 3.2 Latency to *status epilepticus* establishment and mortality rate

All the control and PBS-injected animals developed pilocarpine-induced *status epilepticus* within 155 ± 27.71 and 170 ± 22.36 min, respectively. Regarding the mortality rate, one control and two PBS-injected rats died during the following 72 h. Interestingly, in the GAT1-saporin-injected group, the latency to *status epilepticus* was shorter (140 ± 7.55 min) and the mortality rate higher (three died during the 90-min *status epilepticus* and two within the following 24 h). When analyzing the 192-IgG-saporin-injected group, three animals reached *status epilepticus* (144 ± 24 min; one died the next day) but three did not.

### 3.3 Behavioral evaluations

#### 3.3.1 Effects of cholinergic medial septum lesion on anxiety-related behavior, locomotor activity, and motor coordination

The elevated plus-maze test showed that injecting 192-IgG-saporin into the medial septum had an effect on the anxiety-related behavior of the animals. These rats showed a significantly decreased number of entries [F_(3,28)_ = 3.4; *p* < 0.05] in the open arms (Figure 3A). However, when evaluating the time spent in the open arms, they clearly preferred to stay there, exploring the area [F_(3,28)_ = 5.7; *p* < 0.05] (Figure 3B). This result indicates that damaging cholinergic medial septum neurons can reduce anxiety-related behavior in these animals.

**Figure 3.**
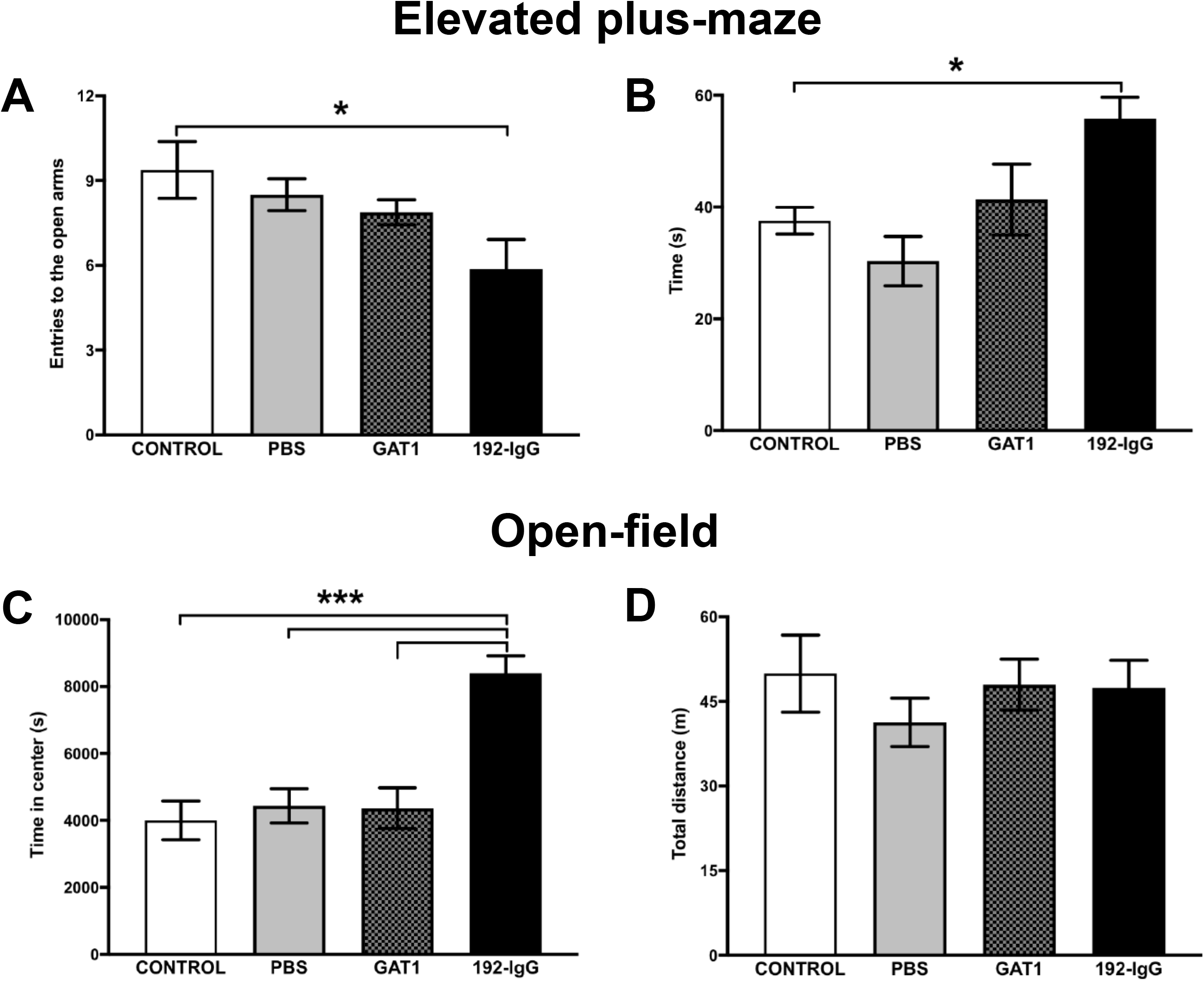
Locomotor activity and anxiety-related behaviors after medial septum lesioning. Reduced anxiety-related behavior is observed in 192-IgG-injected animals, evidenced by the number of entries into the open arms of the elevated plus-maze (**A**) as well as the time spent in such arms (**B**). **C:** supports this notion by evaluating the time that the animals spent in the center of the open field box. **D**: no altered locomotion activity was observed in any of the evaluated groups. Graphs represent the mean ± SEM (\**p* < 0.05, \*\*\**p* < 0.001). PBS, phosphate buffered saline.

The open-field test showed that neither the mechanical lesion nor the injection of saporins into the medial septum had adverse effects on locomotor activity in any of the three groups, which was evident when comparing the four groups’ total distance traveled for 24 h [F_(3,28)_ = 0.5; *p* = 0.67] (Figure 3D). However, when measuring the time spent in the center of the box, only the rats injected with 192-IgG-saporin showed increased values, which can be interpreted as an anxiolytic-like effect in these animals [F_(3,28)_ = 13.85; *p* < 0.001] (Figure 3C).

In the rotarod test, the four groups displayed similar performance throughout the 12 trials, showing motor learning of the test and reaching their optimal plateau on the third day. This demonstrates that coordination was not altered due to the medial septum lesion (Figure 4A). Two-way repeated measures ANOVA values are as follows: Group, F_(3,28)_ = 0.6, *p* = 0.61; Time, F_(5.61,157.2)_ = 21.88, *p* = 0.0001; Interaction, F_(33,297)_ = 0.49, *p* = 0.9911.

**Figure 4.**
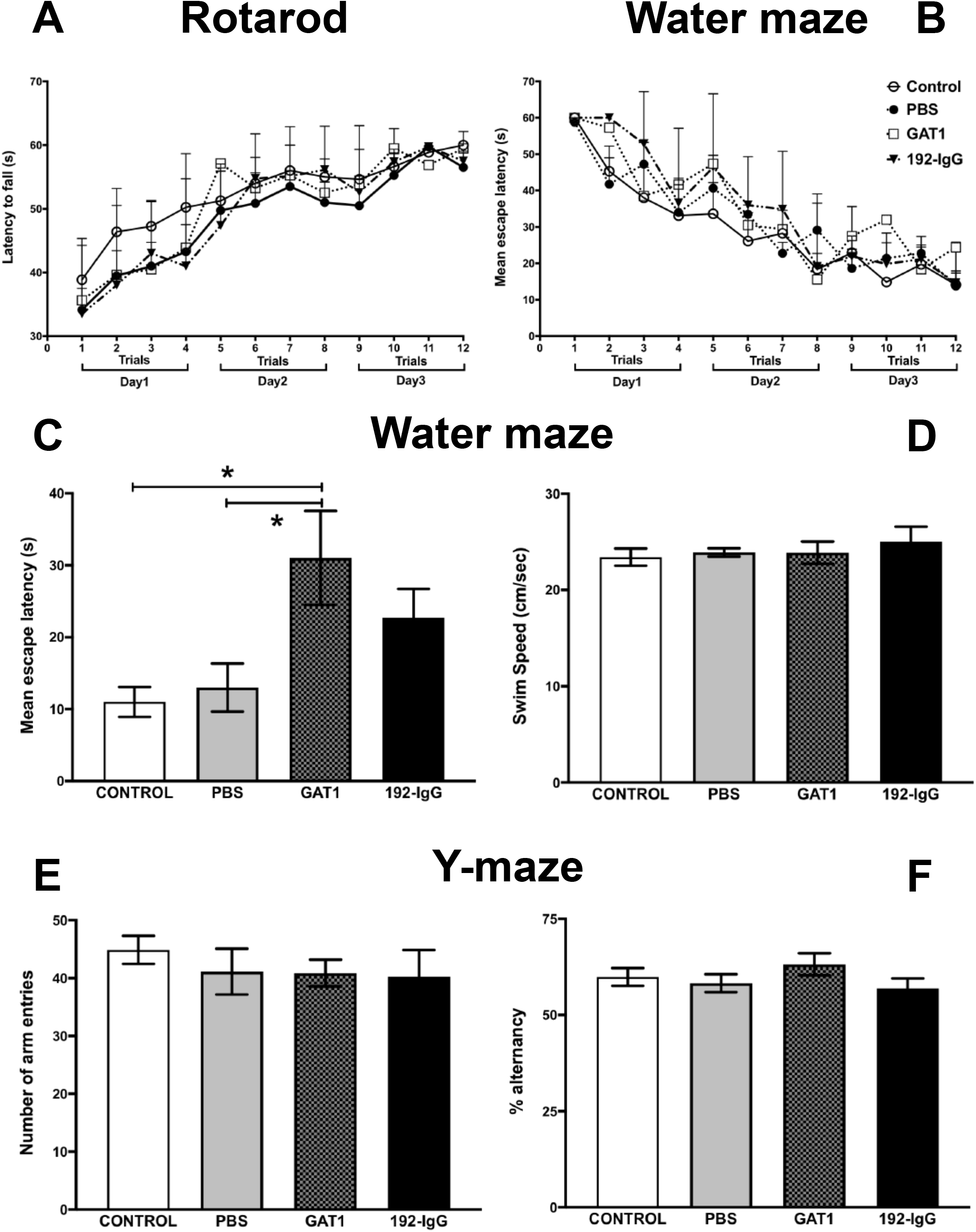
Motor coordination and memory formation. **A**: shows the integrated data of the time spent on the rotating rod for each trial (four trials each day for three consecutive days). No significant differences between groups were observed along the 12 trials. The effect of medial septum lesioning on spatial memory was evaluated by the mean escape latency (mean ± SEM) of animals trained for three days in the Morris water-maze task. **B**: shows no differences between groups during training; therefore, all rats were able to learn the platform’s localization with a similar task acquisition rate. Escape latencies on the fifth day were significantly higher in the GAT1-saporin-injected group (**C**). All the evaluated animals showed no altered swimming speed throughout the experimental procedure (**D**). Working memory was evaluated by the number of arm entries (**E**) and spontaneous alternation behavior (**F**) was measured during a 10-min session in the Y-maze test. Similarities between groups were evident. Bar graphs or temporal courses were analyzed by a one-way ANOVA or a multiple comparisons two-way ANOVA, respectively; both were followed by a *post hoc* Tukey test. \**p* < 0.05 compared to the control and the PBS-injected groups. PBS, phosphate buffered saline.

#### 3.3.2 Spatial memory impairment after GABAergic medial septum lesion

The spatial memory of all the animals was assessed using the Morris water-maze test. A two-way repeated measures ANOVA revealed no differences between groups during training (F_(3,28)_ = 1.12, *p* = 0.35), showing that the animals were able to learn the platform’s localization with a similar task acquisition rate (Figure 4B). On the fifth day, the escape platform was removed and the animals were submitted to a 48 h retention test. Here, the GAT1-saporin-injected animals showed the longest escape latencies when compared to the control and PBS groups (both *p* < 0.05; Figure 4C). In the case of the 192-IgG saporin-injected animals, a similar trend was observed (not significant; Figure 4C). It is important to highlight that swimming speed was not altered in any of the evaluated animals throughout the experimental procedure (Figure 4D).

The spontaneous alternation behavior was used to evaluate working memory in the Y-maze. No significant differences between groups were observed. This can be interpreted as an intact short-term memory since all animals were able to remember which arms they had already visited. Using a one-way ANOVA analysis, we observed no inter-group differences for the number of arm entries [F_(3,28)_ = 0.36; *p* = 0.77] (Figure 4E) nor for spontaneous alternations [F_(3,28)_ = 1.11; *p* = 0.35] (Figure 4F).

### 3.4 *In vivo* DTI analysis

This study focused on the progression of changes after selectively lesioning medial septum GABAergic or cholinergic neuronal populations, as well as differences between groups at each time point (Baseline, 2 weeks and 7 weeks post-injection). ROIs were manually outlined. Figure 5A and 5B shows the delineated regions from both hemispheres. There was a large inter-rater reliability of diffusion metrics derived from all ROIs. Pairwise comparisons of diffusion metrics for bilateral structures did not reveal any asymmetries. Henceforth, the reported values for each ROI represent the average of both hemispheres.

**Figure 5.**
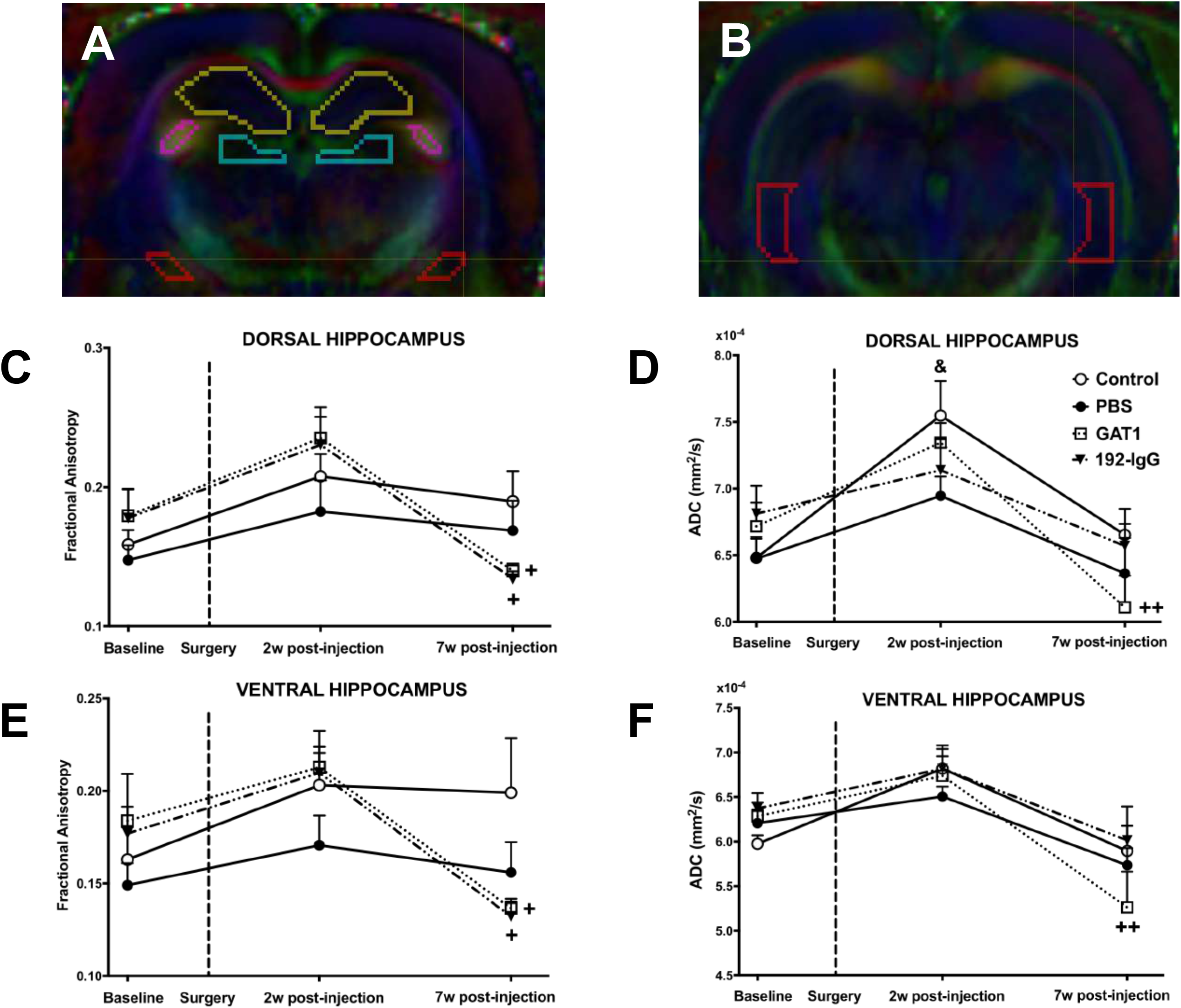
Fractional anisotropy (FA) and apparent diffusion coefficient (ADC) values from dorsal and ventral hippocampi of Control and lesioned rats. **A:** representative coronal images delineating the perimeter of the evaluated brain regions (yellow: dorsal hippocampus; pink: fimbria; blue: dorso-medial thalamus; orange: amygdala). **B**: ventral hippocampus. The graphs show the data obtained using *in vivo* DTI. The dotted line indicates the day of the surgery. The groups injected with the saporins show significantly lower FA values in both hippocampi during the third scanning when compared to the values obtained during the previous scan (**C** and **E**). When comparing ADC values between groups, the GAT1-saporin-injected group shows significant decreases during the last time point (**D** and **F**). + : multiple comparisons two-way ANOVA; * : *post-hoc* between group differences; & : *post-hoc* within-group difference with respect to baseline (one or two symbols for *p* < 0.05 and 0.01, respectively). PBS, phosphate buffered saline; w: weeks.

The dorsal hippocampus showed longitudinal changes in the FA maps of the animals injected with both saporins (2 weeks vs 7 weeks post-injection, ↓44%, *p* < 0.05) (Figure 5C). Regarding ADC, we observed no significant changes between groups. Control (Baseline vs 2 weeks post-injection, ↑17%, *p* < 0.05) and GAT1-saporin-injected animals (2 weeks vs 7 weeks post-injection; ↓17%, *p* < 0.01) showed significant longitudinal changes (Figure 5D).

In the case of the ventral hippocampus, similar longitudinal changes were obtained in the FA maps as those seen in the dorsal hippocampus (2 weeks vs 7 weeks post-injection, ↓38%, *p* < 0.05) (Figure 5E). When evaluating ADC, no significant changes between groups were observed. Only the GAT1-saporin-injected animals (2 weeks vs 7 weeks post-injection; ↓22%, *p* < 0.01) showed significant longitudinal changes (Figure 5F).

Figure 6A shows longitudinal differences in the FA of fimbria in the control (↑11%) and PBS (↑9%) groups when comparing Baseline and 7 weeks post-injection (*p* < 0.05). Regarding ADC, the GAT1-saporin-injected animals showed a significant reduction in the fimbria when compared to the other three groups at 7 weeks post-injection (↓17%, *p* < 0.05) (Figure 6B). Longitudinal changes were observed in control (2 weeks vs 7 weeks post-injection, ↓16%, *p* < 0.05) and GAT1-saporin-injected animals (Baseline vs 7 weeks post-injection, ↓18%, *p* < 0.05, and 2 weeks vs 7 weeks post-injection ↓21%, *p* < 0.01) (Figure 6B).

**Figure 6.**
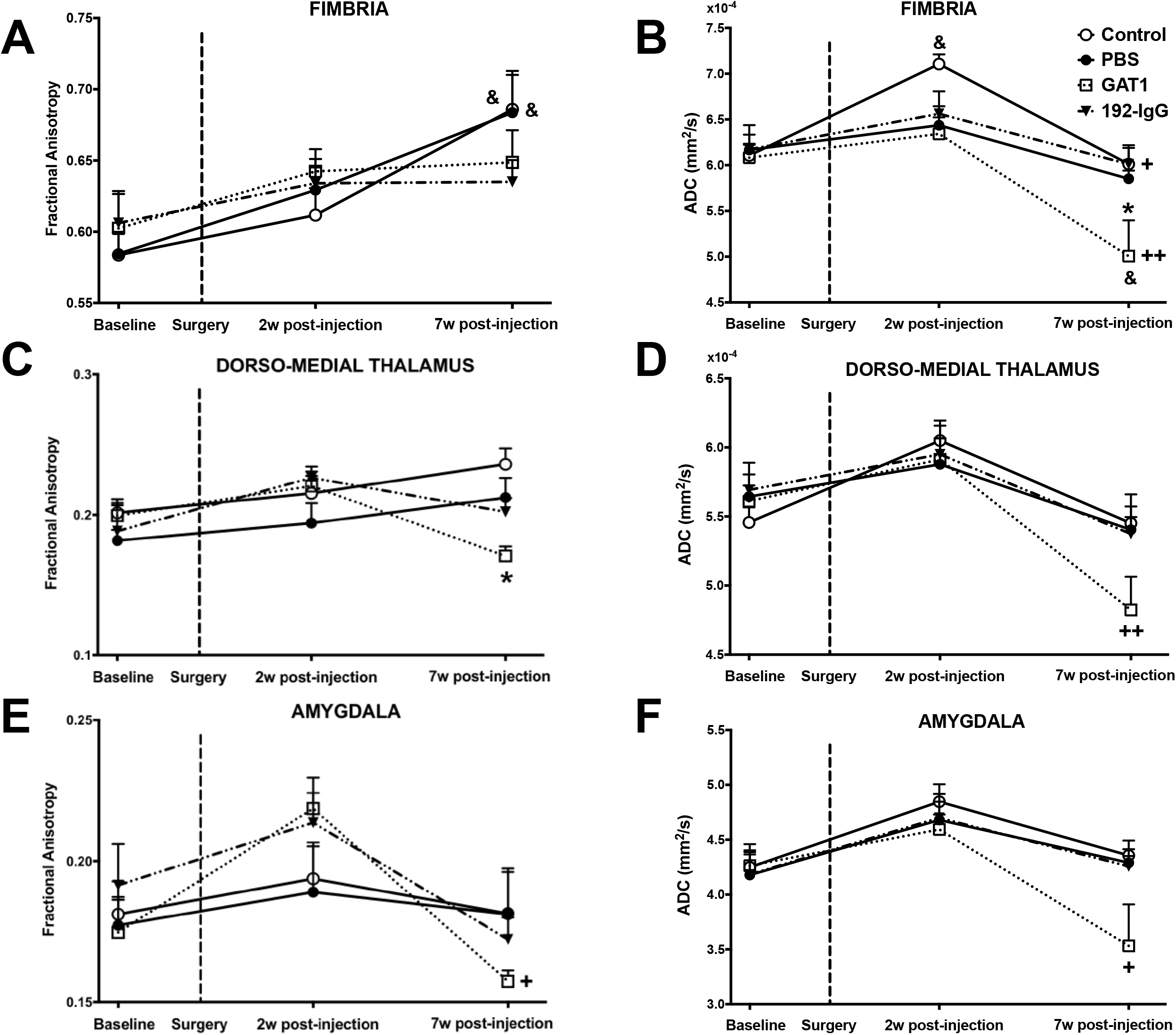
Fractional anisotropy (FA) and apparent diffusion coefficient (ADC) values from fimbria, dorso-medial thalamus and amygdala of Control and lesioned rats. The graphs show the mean and standard error per group. The dotted line indicates the day of the surgery. **A**: in the case of the fimbria, FA values from control and PBS groups show a pattern of increase over time; this effect was not observed when injecting the saporins. **B**: When comparing ADC values, only control animals show a significant increase after 2 weeks post-injection; moreover, the third time-point shows that GAT1-saporin-injected animals are the group with statistically significant changes (longitudinal and between groups). The GAT1-saporin injection into the medial septum also decreased FA and ADC values of the dorso-medial thalamus (**C** and **D**) and amygdala (**E** and **F**). + : multiple comparisons two-way ANOVA; * : *post-hoc* between group differences; & : *post-hoc* within-group difference with respect to baseline (one or two symbols for *p* < 0.05 and 0.01, respectively). PBS, phosphate buffered saline. w: weeks.

The FA maps of the GAT1-saporin-injected animals showed a significant reduction (↓27%; *p* < 0.05) in the dorso-medial thalamus when compared to the control group during the third time point (Figure 6C). Regarding ADC, the same group showed significant longitudinal changes (2 weeks vs 7 weeks post-injection ↓19%, *p* < 0.01) (Figure 6D).

In the case of the amygdala, the FA and ADC values of the GAT1-saporin-injected animals showed significant decreases when comparing 2 weeks and 7 weeks post-injection groups (↓29% and ↓24%, respectively; both *p* < 0.05) (Figure 6E, 6F).

### 3.5 Mossy fiber sprouting and tissue damage

Mossy fiber sprouting in the dorsal hippocampus was evaluated by using Timm staining (Figure 7A-H). There was an evident increase in mossy fiber sprouting in animals injected with the immunotoxin GAT1-saporin (Figure 7C, 7G) compared to the control animals (Figure 7A, 7E). This aberrant sprouting was not present in PBS-(Figure 7B, 7F) or 192-IgG-injected animals (Figure 7D, 7H).

**Figure 7.**
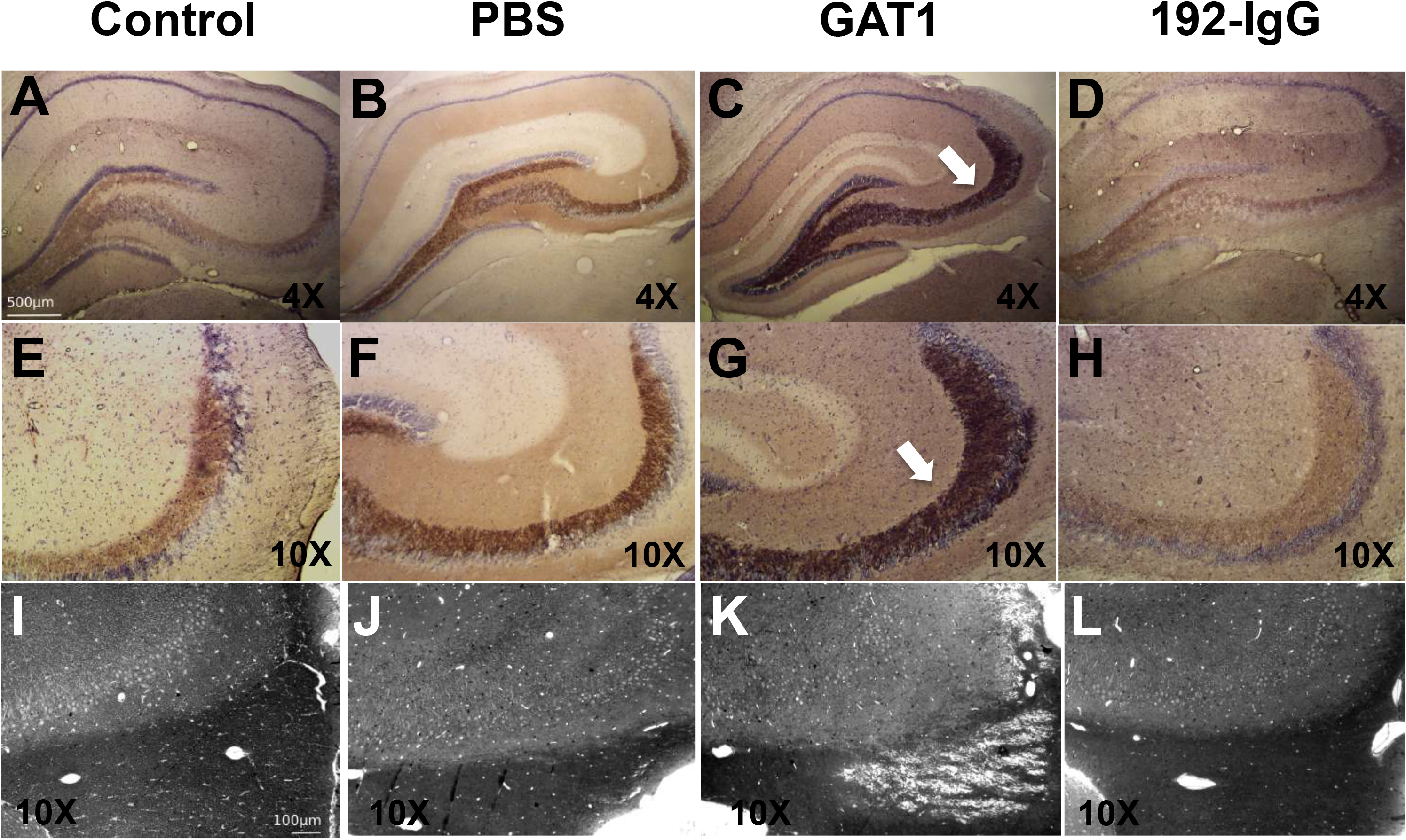
Mossy fiber terminals and tissue damage. First two rows show Timm-stained mossy fibers found in the dorsal hippocampus of normal rats (**A** and **E**) and animals with medial septum injection of PBS (**B** and **F**), GAT1-saporin (**C** and **G**), and 192-IgG-saporin (**D** and **H**). Arrows in **C** and **G** show an aberrant increase of mossy fiber sprouting along CA3 and the dentate gyrus. Mossy fiber sprouting was not observed in the other groups. The third row shows toluidine blue staining microphotographs encompassing the border between the dorsal hippocampus (top) and fimbria (bottom; axonal myelin sheaths stained dark) (**I** – **L**). The injection of GAT1-saporin into the medial septum clearly provokes tissue damage of the fimbria (**K**), an effect not observed in the other groups. PBS, phosphate buffered saline.

Examination of brain tissue slides using toluidine blue staining revealed that animals injected with the immunotoxin GAT1-saporin in the medial septum had tissue damage in the fimbria (Figure 7K). The fimbria was not histologically altered in the animals injected with vehicle (Figure 7J) or the neurotoxin 192-IgG-saporin (Figure 7L).

The observation of Nissl-stained sections revealed considerable damage to the dorsal hippocampus, dorso-medial thalamus and amygdala of GAT1-saporin-injected animals compared to the other three groups. A reduction of pyramidal layer thickness was evident in CA1 and CA3, together with a dispersion pattern of the remanent cells (Figure 8D). In addition, considerable neuronal death was observed in the dorso-medial thalamus (Figure 8H) and amygdala (Figure 8L). The changes in the 192-IgG saporin-injected animals were evident, although not as dramatic as those seen in the GAT1-saporin group.

**Figure 8.**
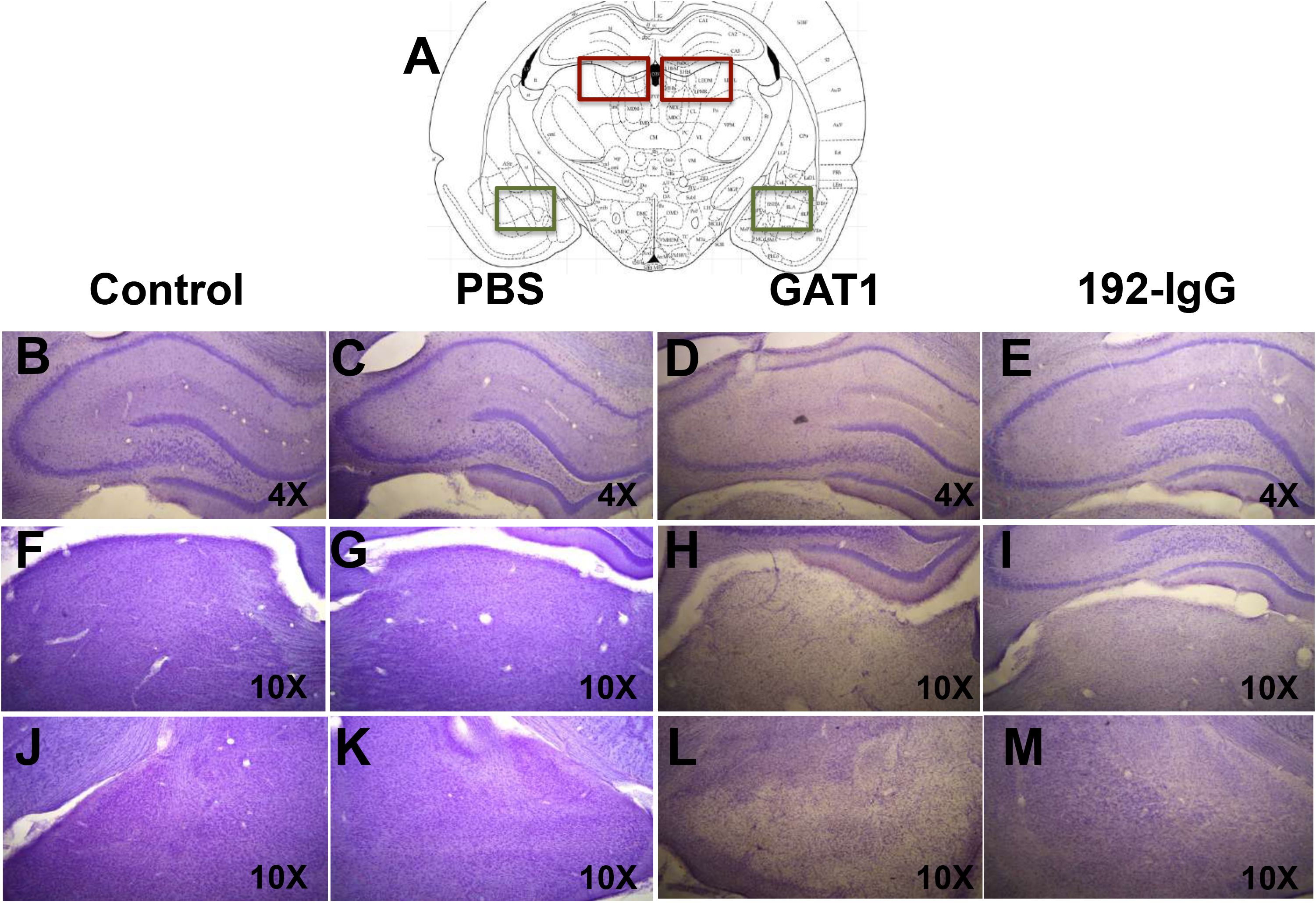
Nissl-stained sections and cell loss. **A**: representative scheme used to frame the evaluated areas (dorso-medial thalamus in red and amygdala in green). First row shows dorsal hippocampus; second row shows dorso-medial thalamus; third row shows amygdala. The injection of GAT1-saporin into the medial septum induces a dramatic cell loss in the three evaluated regions (**D**, **H**, **L**). The injection of 192-IgG-saporin does not cause such an evident impact (**E**, **I**, **M**). No effect is detected after PBS injection (**C**, **G**, **K**). PBS, phosphate buffered saline.

## 4. Discussion

We show for the first time that relatively selective lesions of medial septum cholinergic and GABAergic neurons differentially affect gray and white matter structures relevant to memory formation, anxiety-like behaviors, susceptibility to *status epilepticus* induction, and survival. Parallel to the evolution of tissue damage assessed with DTI, behavioral evaluations were done for one month, revealing no significant differences between control and vehicle-injected rats. However, depending on the task, a dissociation among the saporin-injected animals was observed. Histologically, GAT1-saporin significantly damaged fimbria and induced neuronal loss in the dorsal hippocampus, dorso-medial thalamus and amygdala. A striking difference between groups was the presence of aberrant mossy fiber sprouting after lesioning the septal GABAergic neuronal population, a common histopathological feature in patients with TLE and reproduced in animal models. All these changes induced in normal animals by medial septum alterations suggest the establishment of a hyper-excitable network likely predisposing to epileptiform activity.

The medial septum is an interconnected brain region encompassing two major populations of neurons, the parvalbumin-containing GABAergic cells (Freund, 1989) and the ChAT-immunoreactive cholinergic neurons (Rye et al., 1984). Having different hippocampal targets: the former innervate GABAergic hippocampal interneurons (Freund and Antal, 1988), which then synapse onto pyramidal cells (Toth et al., 1997); the latter contact pyramidal and dentate granule cells, and inhibitory interneurons (Frotscher and Leranth, 1985). Thus, the medial septum constitutes a source of hippocampal innervation via the fimbria-fornix, where the differential target selectivity of each septal neuronal population may have a different role in the regulation of hippocampal electrical activity (Hangya et al., 2009; Levesque and Avoli, 2018). This has increased the medial septum’s relevance as a potential modulator of the septo-hippocampal interface in memory formation and the pathophysiology of neurological diseases such as epilepsy.

192-IgG-saporin has been used as a toxic agent of the cholinergic neurons (Pioro and Cuello, 1990; Wiley et al., 1991). Previous studies, including ours, have reported that the injection of 192-IgG-saporin into the medial septum reduces the cholinergic markers within seven days, with a slight decrease in GABAergic septo-hippocampal (parvalbumin-immunoreactive) neurons (Craig et al., 2008). Regarding hippocampal-dependent working memory, we found no differences, in line with previous reports (Chappell et al., 1998; Pang et al., 2001; Kirby and Rawlins, 2003). However, when evaluating spatial memory formation, a slight but not significant impairment was observed (Berger-Sweeney et al., 1994; Lecourtier et al., 2011; Dashniani et al., 2020). We discard the possibility that such an effect can be attributable to motor coordination deficits in the animals due to a mistaken injection site (Lehmann et al., 2000) since no significant differences between groups were observed in the swimming speed or during the open field (total distance parameter) and rotarod tests. Therefore, our results suggest that 1) the septo-hippocampal cholinergic input is involved in hippocampal-dependent memory processing, but other neuronal systems are also required for this to occur (McIntyre et al., 2002); 2) the time at which behavioral testing is conducted after lesioning is crucial (Paban et al., 2005); and 3) the medial septum-induced lesions compromise the integrity of other brain regions such as the fimbria-fornix, dorsal hippocampus and dorso-medial thalamus, an effect that increases the chance of impairing the animals’ spatial memory process (Markowska et al., 1989; Compton, 2004; Taube, 2007; Miyoshi et al., 2012). Our results can shed some light on a previously raised hypothesis related to Alzheimer’s disease development and age-related memory deficits (Mesulam, 1986; Gallagher and Colombo, 1995; Paban et al., 2005; Schliebs and Arendt, 2011; Palop and Mucke, 2016), where the authors suggest a specific loss of cholinergic innervation of the hippocampal formation prior to cell death, axonal degeneration and plaque formation.

The septo-hippocampal cholinergic system has also been implicated in the maintenance of innate anxiety in rodents (Zarrindast et al., 2013; Zhang et al., 2017). We did not observe significant changes in anxiety-like behaviors in PBS- and GAT1-saporin-treated rats. In contrast, when injecting 192-IgG-saporin, our animals showed reduced anxiety-like behaviors. Effect previously described following lesions of the medial septum cholinergic neurons through other techniques (Pesold and Treit, 1992, 1994; Pizzo et al., 2002; Zhang et al., 2017; Jiang et al., 2018). In our case, we discard impaired locomotor activity as a potential factor affecting the animals’ behavior. Then, such changes can possibly be attributable to the microstructural alterations of the ventral hippocampus and amygdala seen with *in vivo* DTI, mainly based on: 1) the way the medial septum connects with the hippocampus and how the afferent and efferent connections of this structure vary along its dorso-ventral axis; specifically, the ventral hippocampus projects to the prefrontal cortex, hypothalamus, and amygdala (Swanson and Cowan, 1977; Fanselow and Dong, 2010); and 2) previous reports describing reduced anxiety-like behavior after ventral hippocampal (Bannerman et al., 1999, 2004) or amygdala lesions (Ranjbar et al., 2017). Overall, our results provide evidence that lesioning the medial septum cholinergic neurons can reduce anxiety-like behavior in the animals without significantly affecting their working and spatial memory.

We also administered a GABAergic immunotoxin into the medial septum to examine the importance of this neuronal population in various behavioral processes. As previously reported, GAT1-saporin significantly reduced the number of parvalbumin-immunoreactive GABAergic neurons within seven days without considerably altering the number of the cholinergic ones (Pang et al., 2011; Köppen et al, 2013; Roland et al., 2014a). The similarity also lies in the fact that animals with this type of damage showed impaired spatial memory (Lecourtier et al., 2011; Pang et al., 2011) as well as intact working memory when subjected to short retention intervals (Pang et al., 2001; Roland et al., 2014a). Effects possibly due to: 1) the damage generated on other non-GABAergic neurons located in the medial septum, such as the cholinergic cells (Pang et al., 2011); 2) the impact on the ventral hippocampus acetylcholine efflux (Amaral and Kurz, 1985; Roland et al., 2014a); 3) the disruption of the hippocampal theta rhythm function (Hasselmo, 2005; Yoder and Pang, 2005); and 4) the lesion magnitude at the time at which behavioral testing is conducted (Paban et al, 2005), as evidenced by the *in vivo* DTI data, the tissue damage at the border between fimbria and dorsal hippocampus, and the mossy fiber sprouting detected in CA3. As previously mentioned, unlikely that spatial memory impairment is due to the motor coordination deficits, which are sometimes attributable to a mistaken injection site.

A striking result is the presence of aberrant mossy fiber sprouting in the dorsal hippocampus of GAT1-saporin-injected rats. We can suggest, based on the studies by Hannesson et al. (1997) and Mohapel et al. (1997), that after a bilateral transection of the fimbria/fornix, the septo-hippocampal GABAergic neuronal pathway is possibly involved in the aberrant sprouting of the dentate gyrus mossy fibers. The factors involved in the generation of mossy fiber sprouting are still not completely understood, and there is considerable debate about whether they are a cause or consequence of seizures (Acsády et al, 1998; Buckmaster et al., 2002; Cavazos et al., 2003; Kim et al., 2003; Koyama et al., 2004; Evstratova and Toth, 2014). Researchers have used electron microscopy and the histochemical Timm sulfide silver method to visualize axonal projections and terminations of the dentate mossy fibers under normal and experimental conditions while also evaluating lesion-induced changes in neuronal connectivity (Zimmer, 1973; Danscher, 1981; Laurberg and Zimmer, 1981; Frotscher and Zimmer, 1983). In epilepsy-related conditions, the dentate mossy fibers branch out and aberrantly project to diverse molecular layers, including reverse projections that form excitatory synapses on the granule cell dendrites, a situation hypothesized to play a crucial role in the generation of seizure activity (Buckmaster et al., 2002; Cavazos et al., 2003). Abundant sprouting has been reported in patients with drug-resistant TLE (Mathern et al., 1995; Proper et al., 2000; Blümcke et al., 2012, 2013; Schmeiser et al., 2017), the most common form of focal-onset epilepsy (Tellez-Zenteno and Hernandez-Ronquillo, 2012). Mossy fiber sprouting has also been described in epilepsy animal models (Cavazos et al., 1991; Levesque et al., 2016). However, a big debate exists regarding the role that mossy fiber sprouting plays in epileptogenesis and chronic epilepsy. Some authors argue that it is compensatory (Sloviter et al., 2006; Schmeiser et al., 2017) while others claim that it is epileptogenic (Feng et al., 2003; Hendricks et al., 2019). In our case, the GAT1-saporin-injected rats did not exhibit altered behavior typical of an epileptogenic process, despite this histopathological phenomenon. However, mossy fiber sprouting can be considered a risk factor capable of influencing not only seizure susceptibility but also mortality rate. Recent studies in TLE models suggest that targeting specific neuronal populations within the septo-hippocampal pathway can be a promising tool to understand epileptogenesis and find new treatment strategies to block seizures (Bao et al., 2020; Wang et al., 2020; Hristova et al., 2021; Velasquez et al., 2023).

In this study, we used DTI to infer tissue characteristics (Basser et al., 1994). Demonstrating for the first time the tissue abnormalities that occur over time following the injection of vehicle or saporins into the medial septum. Both toxins induced significant changes 7 weeks post-injection in gray and white matter structures relevant to seizure initiation and propagation in TLE. Interestingly, such a phenomenon occurred, but none of the animals showed behaviors typical of an epileptogenic process.

Regarding the dorsal hippocampus, diffusion abnormalities are more pronounced after GAT1-saporin injection and can be attributable to large-scale neuronal loss and mossy fiber sprouting. Similar findings to those reported in animal models of epilepsy (Laitinen et al., 2010; Parekh et al., 2010; Luna-Munguia et al., 2021), where are considered as crucial factors for generating aberrant hyperexcitable networks (Scharfman et al., 2000). Other factors should also be taken into account, including the reorganization of myelinated axons (Laitinen et al., 2010; Sierra et al., 2015), the activation of inflammatory cells (Pernot et al., 2011), and the dendritic arborization of granule cells (Sierra et al., 2015). However, in contrast to previous reports showing increases at chronic time points of epileptogenesis (Salo et al., 2017; Luna-Munguia et al., 2021), our results reveal a significant decrease in FA values at 7 weeks post-GAT1-saporin or 192-IgG-saporin injection.

In saporins-injected rats, the fimbria did not show the upward trend in FA seen in the control and vehicle-injected groups after 7 weeks post-injection. Diverse DTI studies involving patients with TLE and animal models of chronic TLE have reported bilateral diffusion abnormalities of the fimbria and subsequent significantly reduced FA values (Concha et al., 2005b, 2007; Parekh et al., 2010; van Eijsden et al., 2011; Otte et al., 2012; Luna-Munguia et al., 2021). Altered diffusion metrics of the fimbria reported in patients with TLE have been interpreted as a sensitive way to detect down-stream effects of hippocampal cell loss (i.e., degeneration of hippocampal efferents) (George and Griffin, 1994; Concha et al., 2005a, 2010). Although DTI cannot distinguish between afferent and efferent fiber populations within the bi-directional fimbria, the direct and selective lesions of the medial septum in our current study support the notion of hippocampal afferent degeneration. While analyzing fimbria ADC values longitudinally, the GAT1-saporin-injected animals showed the largest abnormalities in this structure after 7 weeks post-injection. This dramatic reduction can be attributable to a restricted motion of intra- and extracellular water compartments due to several factors (Gass et al., 2001) and to macrophage, microglial and astrocytic proliferation (Wall et al., 2000). Similar effects have been reported clinically (Szabo et al., 2005) or in experimental *status epilepticus* models (Fabene et al., 2003; Engelhorn et al., 2007). While the propensity to develop spontaneous seizures was not evident in our experimental animals within the time frame studied, disruption of the septo-hippocampal network may induce long-term effects that favor epileptogenesis and explain the diffusion abnormalities of the fimbria seen early in the course of TLE (van Eijsden et al., 2011) and their association with the development of chronic seizures in rodents (Parekh et al., 2010).

Marked FA changes were also observed within the dorso-medial thalamus in both saporin-injected groups. Relevant considering that thalamus is an extra-limbic structure that has excitatory influence over the hippocampus (Bertram and Zhang, 1999), exerting a regulatory role in the initiation and propagation of limbic seizures (Cassidy and Gale, 1998). In contrast, Parekh and co-authors (2010) did not report any alteration during the latent period, a result that complicates the association between thalamic changes and the onset of spontaneous seizures. Regarding the amygdala, they suggest that *status epilepticus*-induced damage to this region may be irreversible and able to trigger the onset of spontaneous seizures. In our case, GAT1-saporin-injected rats showed a significant drop in FA and ADC values at 7 weeks post-injection. As in previous reports, this coincides with the evident decrease in neuronal density (Wall et al., 2000).

We provide evidence of time-dependent diffusion changes in gray and white matter after injecting both saporins. The characterization of tissue damage over time provides new evidence of the impact that medial septum lesioning has on the integrity of limbic regions and how these changes can represent a precursor for behavioral deficits or epilepsy development. Our findings support the idea that modulation of the medial septum can be a potential target to enhance cognition or reduce seizure frequency, improving the patients’ quality of life.

## Author Contributions

HL-M, DG-M, and LC contributed to conception and design. HL-M, DG-M, and AG-C performed the surgeries, behavioral tests, and imaging. HL-M, DC, MR, ER, and PV performed the histological procedures. HL-M, DG-M, AG-C, DC, FH-F, KF-G, BYB, and LC worked on the analysis and interpretation of data; HL-M, DG-M, and LC were involved in drafting the manuscript and revising it critically. All authors revised and read the manuscript and approved the submitted version.

